# Pilot Analysis of Genetic Effects on Personality Test Scores with AI: ABO Blood Type in Japan

**DOI:** 10.1101/2023.01.15.524099

**Authors:** Masayuki Kanazawa

## Abstract

It is estimated that genetic factors are responsible for approximately 30-60% of an individual’s personality. However, statistical analyses have yet to demonstrate a significant and consistent relationship between personality tests and genetic factors. Conversely, a significant portion of individuals in Japan, South Korea, and Taiwan believe that there is a linkage between ABO blood type, which is determined genetically, and personality. This pilot study analyzed data from a large-scale survey (N=2,887) using a combination of traditional statistical methods and AI to examine this relationship. The results indicated the relationship between ABO blood type and self-reported personality traits on several single-question items, in accordance with expected outcomes. These findings suggest that this relationship may extend beyond ABO blood type and personality to encompass other inherited characteristics. The influence of the level of interest on personality and was also discussed.

## 1. Introduction

### 1.1. Influence of Heredity Based on Twin Studies

The estimated percentage of heredity’s impact on personality ranges from 30-60% according to twin studies and other investigations [1–4]. There exist two distinct types of twins: identical and fraternal. Identical twins are formed from a fertilized egg that splits during multiplication, resulting in both individuals having identical DNA. Conversely, fraternal twins are formed from the fertilization of two separate eggs by different sperm, resulting in them having distinct DNA profiles. In reality, it is observed that identical twins often have the same facial features, hair color, and eye color, while fraternal twins do not typically exhibit such similarities.

In light of these findings, the influence of heredity on psychological and physical characteristics has been evaluated through statistical analysis of the differences between identical and fraternal twins. To date, a plethora of twin pairs have been studied through the use of personality tests such as the Big Five. The outcomes of these studies indicated that differences between identical twins were minimal in comparison to the differences between fraternal twins, suggesting that psychological and constitutional traits are partially determined by DNA. These twin studies have been conducted globally with comparable results, and the data generated is highly reproducible. However, it should be noted that these surveys primarily focused on standard families in developed countries, and therefore did not encompass the more severe conditions, such as malnutrition during times of war.

In the 1950s, it was discovered that heredity is regulated by a biomolecule with a double helical structure known as DNA (deoxyribonucleic acid). DNA encodes genetic information through the sequence of 4 base classes, comprising billions of bases in humans. Due to the recent rapid advancements in the life sciences, the International Human Genome Sequencing Consortium announced in 2003 that the complete DNA sequence, or whole genome, had been decoded. The subsequent 1000 Genomes Project, initiated in 2008, was the first among many genome sequencing projects that have been launched globally, yielding a plethora of solid results. These developments have enabled the direct analysis of the relationship between genomes and personality and abilities through mathematical methods, which was not possible using traditional methods.

### 1.2. Genome-wide Association Analysis Studies

A prevalent methodology employed in genetics research is GWAS (genome-wide association analysis studies), which aims to identify genetic variations associated with specific diseases or characteristics of an individual’s constitution. This is accomplished by collecting and comprehensively analyzing the genomic information of a large population to identify small variations in genetic information known as SNPs (single nucleotide polymorphisms). Each analysis examines hundreds to thousands of SNPs simultaneously and compares the differences in genetic information found between populations that possess a specific trait and those that do not, in order to identify the SNPs that appear more frequently in relation to that trait.

However, even when SNPs are identified through GWAS, the genetic contribution derived from them is typically significantly smaller than that estimated through twin studies. This phenomenon, known as missing or hidden heritability, has garnered significant attention as GWAS is increasingly utilized.

A notable example of this is provided by Lo et al [5]. In this study, the researchers sought to correlate data on all SNPs in the human genome of over 260,000 individuals with the Big Five personality test, the most widely used personality test in current psychology. As a result, 6 genes were identified as being associated with personality, however, the coefficient of determination *R^2^*, which indicates the degree of influence, was less than 0.4% for all of them, an extremely small value that would typically be considered an error. This value is much smaller than the 30-60% values obtained in twin studies.

There are several potential explanations for such extremely small heritability estimates, including 1) interactions between genes (nonlinear interactions), 2) genetic and environmental interactions, and 3) the challenge of detecting and analyzing SNPs with current molecular genetics techniques. This is also important in approaching an essential understanding of genetic phenomena. Recent studies have shown that over 1,200 genes are involved in height [6] and 124 genes in hair color [7].

### 1.3. Blood Type Personality Studies

Conversely, some studies have arrived at the opposite conclusion from the aforementioned explanation, that the effects of individual genes are minuscule. This is exemplified by studies on the correlation between blood type and personality. A significant number of studies on this topic have been published in East Asia, particularly in Japan.

Approximately half or more of the population in Japan, South Korea, and Taiwan contend that the relationship between genetically determined ABO blood type and personality is credible [8–13]. However, there is still a lack of scientific consensus on the relationship between blood type and personality. What past research findings have in common is that the Big Five personality tests, which aggregate multiple questions from 5 to several hundred into 5 factors, have shown little or no consistent differences [12–15]. In contrast, many studies have found statistically consistent and significant differences when analyzing each single question item that is considered to be a “blood type trait.” This can be successfully explained by considering that blood type affects only a very limited number of question items that comprise the personality factors, which would explain the previously inconsistent results. This is because each personality factor is a statistical integration of many question items, and the differences in the few question items that differed by blood type are expected to be reduced to negligible levels. This is similar to the previously mentioned mechanism by which genes that affect height and hair work [6–7].

Since the year 2000, several studies have been published to explore the relationship based on genetic factors such as linkage disequilibrium [16–17], and another report found a relationship between genotype and the TCI Personality Test, as predicted by blood type [18]. In 2022, several studies on ABO blood type and intestinal and intestinal microflora were published [19–21]. According to these reports, differences in the abundance of bacteria have been shown to have a causal relationship with depression and mental health.

### 1.4. Strategy of This Study

As previously stated, hundreds and thousands of genes are involved in macroscopic data such as height and hair color [6–7]. If one assumes that the properties of the Big Five factors are similar to those of height, it is at least theoretically possible to assume that these two mechanisms are identical. As previously noted, the Big Five personality test ultimately narrows the thousands of potential question items down to no more than a few hundred. This is because question items with relatively small differences are omitted. If genes act on each item separately and each effect is small, then many question items that differ by blood type would be omitted. Accordingly, the final version of the personality test resulted in little or small differences in personality factors. This aligns with the fact that previous studies have not yielded consistent differences, and is very similar to the behavior of a number of genes that influence height and hair.

While testing this hypothesis would require a very large amount of personality and genomic data, “blood type” would significantly improve the situation, as more than 90% of Japanese people know their blood type. If 100% accuracy is not required, an Internet survey is sufficient to complete the task. As for the Big Five personality tests, the Big Five Scales (BFS), standardized in Japan, includes blood type traits in some of the 60 items that make up the 5 personality factors [22].

In recent years, there has been a proliferation of scales that aim to measure psychological constructs with a minimal number of items, such as scales to measure subjective well-being [23] and self-esteem [24] with a single item. These scales have been widely adopted in various studies. Similarly, in the realm of the Big Five personality test, brief measures such as 5-item and 10-item inventories have been developed. For instance, Gosling, Rentfrow, and Swann created the Ten Item Personality Inventory (TIPI), which measures the 5 factors of the Big Five in 10 items, using a 7-point scale [25]. Furthermore, Oshio, Abe, and Cutrone developed a Japanese version of the Ten Item Personality Inventory (TIPI-J) [26].

If personality tests are capable of measuring differences in personality based on blood type, certain question items within the test should exhibit variations based on blood type. Thus, the following methodology will be employed in this study. In the Big Five personality test, the number of question items varies from five to several hundred. Therefore, utilizing data obtained from the “same sample,” we will compare the results of the aforementioned 10-item test [26] with those of a test comprising a larger number of items. If the differences of blood type traits are small, then many of them are omitted, a test with more items should be able to detect blood type differences more effectively. Blood type traits are typically described by adjectives, such as “nervous.” Thus, a Big Five personality test whose question items are adjectives and which includes blood type traits should be employed for comparison. One such Japanese-language personality test that satisfies this condition is the aforementioned BFS; this test includes 3 blood type traits in all 60 question items [22]. Therefore, we will investigate whether blood type differences appear not only in these 3 question items, but also in the personality factors that include these items.

### 1.5. Analysis Using AI

Although personality is known to interact with a complex interplay of genetic factors such as gender and age, previous findings have shown that the interaction between these factors is not always linear [27–29]. Most previous psychological studies have assumed that the influence of blood type is constant, regardless of gender, age, and other factors [9–15]. Statistical methods used in designing personality tests using questionnaire methods, such as principal component analysis and correlation analysis, fundamentally assume linear relationships. Therefore, there is no theoretical guarantee that they can accurately analyze data that are not always linear in reality.

These issues may be addressed through the use of artificial intelligence (AI or machine learning). This is because AI can theoretically handle such nonlinear models. Therefore, in this study, we decided to pilot the use of AI to predict blood types from the results obtained from the question items, as well as traditional ANOVA. Since this prediction is typical of “supervised learning,” there is no need to prepare a special mathematical model; current AI cloud systems can handle this task. Assuming the aforementioned mechanism, i.e., that genes act on each question item separately and that the effect is small and omitted for personality tests with fewer items, the prediction of participants’ blood types using blood type traits is more accurate than the prediction using the 5 factors of the Big Five personality test.

## 2. Methods

### 2.1. Participants and Question Items

Given that the implementation of AI necessitates a substantial amount of data, we elected to utilize crowdsourcing as a means of data collection for this study. A sample size of 3,200 individuals was selected for the year 2022, and participants were administered two commonly utilized personality tests, the BFS and the TIPI-J. The sample was stratified by age, with each decade represented, and evenly distributed between males and females. Participants were asked to rate each question item on a scale of 1 to 7, with greater numbers indicating a stronger agreement with the statement. Additionally, participants were asked to rate their belief and knowledge of blood type personality traits on a score of 1 to 4, with greater numbers indicating a greater belief or knowledge. Self-reported blood type was utilized, as a majority of Japanese individuals are aware of their blood type. Out of 3,200 participants, 161, or 5% of the total, were unaware of their blood type. Of the remaining 3,039 participants, 2,887 were used in the final analysis, after excluding any apparently invalid samples, such as those who provided identical responses to the first 20 sequential questions of the personality test. Lastly, an additional item was included to assess participants’ interest in their own and/or others personalities, and participants were asked to rate their interest on a scale of 1 to 7, with greater numbers indicating a greater interest.

In order to circumvent ethical concerns, the blood type trait question items were extracted from previously published literature and academic studies [9–10]. As previously stated, the BFS includes 3 items related to blood type traits, an additional 5 items were added, resulting in a total of 8 items; 2 items for each of the 4 blood types. These 5 question items were subsequently reviewed by a Japanese crowdsourcing company that holds the Japan’s Privacy Mark (JIS Q15001) certification, and were confirmed to not present any ethical issues. This company provides anonymous data to its clients and informed consent was obtained from participants in advance. This study also obtained prior approval from the institutional Ethics Review Committee. The statistical software used for ANOVA was jamovi 2.3.18, and the significance level was set at 0.05. The blood type distribution was 1,091 type As, 632 type Bs, 862 type Os, and 302 type ABs, which were consistent with the Japanese population average [30]. Effect sizes were also calculated [31]. For multiple comparisons, the Holm’s method was utilized.

### 2.2. Analytical Strategy

We tried to follow those methods of psychological personality testing, and deliberately selected the most suitable traits (Table 1) that would certainly produce the differences: images of traits were consistent to the preceding academic studies, they showed large differences in the academic studies and their means were close to 50%, they did not show extreme values, and they were consistent to the preceding surveys of other studies.

**Table 1.** Blood Types and Its Major Personality Traits

| Blood Type | Personality |
| --- | --- |
| **A** | Meticulous* (BFS: Conscientiousness), Nervous* (BFS: Neuroticism) |
| **B** | Self-paced, Self-centered* (BFS: Agreeableness) |
| **O** | Big-hearted, Laid back |
| **AB** | Hard to be understood, Dual personality |

\* Three items included in the BFS.

We utilized traditional ANOVA, and the Microsoft Azure Machine Learning, a cloud AI system. All question items, including gender and age, were used as training data for prediction targeting for the blood type. All data was divided into training and predicting. This was done automatically by the system. In these cases, since the sample sizes of the AI training data were relatively small (this means that the prediction errors might become larger if we used the raw data of 1-year increment of age), a dummy variable of 10-year increments was used [20s = 2, 30s = 3, 40s = 4, 50s = 5].

AUC was used to evaluate the predictions; AUC is a metric used to evaluate binary classification. It has a value of 0.5 if machine learning does not work (the prediction is the same as a random result), and a value of 1 for a perfectly correct prediction. For multiple classifications with 4 values, such as blood type, the weighted AUC is used instead of the standard AUC. Similarly, the value of the weighted AUC is 0.5 if the prediction is the same as the random result and 1 if it is a perfect prediction.

## 3. Results

### 3.1. Analysis 1: ANOVA for the Big Five Personality Factors and Blood Type Traits

As for the BFS, only one factor, Conscientiousness, was statistically significant at *p* < 0.05, and after the Holm’s correction. The magnitude of *η^2^* was at most 0.010, and thus the effect size was small (Table 2). As for the TIPI-J, no factor was significant (Table 3).

**Table 2.** ANOVA for the Big Five Personality Factors of the BFS and Blood Type

| Factors | Average Scores by Blood Type | | | | *F* | $\eta^2$ | *p* |
| --- | --- | --- | --- | --- | --- | --- | --- |
|  | A<br>(N=1091) | B<br>(N=632) | O<br>(N=862) | AB<br>(N=302) |  |  |  |
| **BFS-N (Neuroticism)** | 4.298 | 4.308 | 4.311 | 4.273 | 0.372 | 0.000 | 0.773 |
| **BFS-E (Extraversion)** | 3.870 | 3.822 | 3.846 | 3.871 | 0.395 | 0.000 | 0.756 |
| **BFS-O (Openness)** | 3.759 | 3.755 | 3.789 | 3.744 | 0.179 | 0.000 | 0.910 |
| **BFS-A (Agreeableness)** | 4.321 | 4.262 | 4.279 | 4.237 | 1.128 | 0.001 | 0.278 |
| **BFS-C (Conscientiousness)** | 3.689 | 3.876 | 3.873 | 3.819 | 10.141 | 0.010 | **< .001\*** |

Note. $p < 0.05$ are highlighted in bold. \* $p < 0.05$ after the Holm's correction.

**Table 3.** ANOVA for the Big Five Personality Factors of the TIPI-J and Blood Type

| Factors | Average Scores by Blood Type | | | | *F* | $\eta^2$ | *p* |
| --- | --- | --- | --- | --- | --- | --- | --- |
|  | A<br>(N=1091) | B<br>(N=632) | O<br>(N=862) | AB<br>(N=302) |  |  |  |
| **Neuroticism** | 4.159 | 4.186 | 4.148 | 4.113 | 0.283 | 0.000 | 0.838 |
| **Extraversion** | 3.544 | 3.514 | 3.544 | 3.599 | 0.338 | 0.000 | 0.798 |
| **Openness** | 3.726 | 3.765 | 3.680 | 3.831 | 1.478 | 0.002 | 0.218 |
| **Agreeableness** | 4.689 | 4.646 | 4.586 | 4.680 | 1.460 | 0.002 | 0.224 |
| **Conscientiousness** | 3.946 | 3.911 | 3.867 | 3.871 | 0.771 | 0.001 | 0.510 |

As for the blood type traits, all items were statistically significant at *p* < 0.05, and after the Holm’s correction. The magnitude of *η^2^* was at most 0.026, and thus the effect size was small (Table 4). In 7 of the 8 items, with the exception of the item “self-centered,” the blood type with the highest score matched the respondent’s blood type traits.

**Table 4.** ANOVA for the Blood Type Traits

| Traits (Blood Type, BFS) | Scores by Blood Type | | | | *F* | $\eta^2$ | *p* |
| --- | --- | --- | --- | --- | --- | --- | --- |
|  | A<br>(N=1091) | B<br>(N=632) | O<br>(N=862) | AB<br>(N=302) |  |  |  |
| Nervous (A, N) | **4.271** | 4.079 | 4.200 | 4.017 | 3.880 | 0.004 | **0.009\*** |
| Meticulous (A, C) | **4.335** | 4.006 | 4.044 | 4.060 | 11.240 | 0.012 | **< .001\*** |
| Self-centered (B, A) | 3.681 | 3.938 | 3.823 | 3.967 | 6.630 | 0.007 | **< .001\*** |
| Hard to be understood (AB, N/A) | 3.897 | 4.112 | 4.085 | **4.182** | 5.880 | 0.006 | **< .001\*** |
| Self-paced (B, N/A) | 4.487 | **4.812** | 4.587 | 4.546 | 8.021 | 0.008 | **< .001\*** |
| Dual personality (AB, N/A) | 2.929 | 3.082 | 3.060 | **3.772** | 25.186 | 0.026 | **< .001\*** |
| Big-hearted (O, N/A) | 4.097 | 4.103 | **4.289** | 4.007 | 5.621 | 0.006 | **< .001\*** |
| Laid back (O, N/A) | 3.899 | 4.084 | **4.462** | 4.053 | 25.510 | 0.026 | **< .001\*** |

Note. Highest scores that match with blood type traits and $p < 0.05$ are highlighted in bold. \* $p < 0.05$ after the Holm's correction.

On the question item “Do you think blood type and personality are related?”, 9.0% of the respondents answered “related a lot,” 47.9% answered “related somewhat,” 25.8% answered “not related,” and 17.3%answered “I don’t know.” On question item “Do you know the traits and compatibilities of blood types?”, 4.5% answered “I know a lot,” 44.0% answered “I know some,” 35.9% answered “I know a little,” and15.6% answered “I don’t know at all.”

### 3.2. Analysis 2: Blood Type Predictions Using AI

AUCs which indicate the accuracy of predictions were, in descending order, 0.593 for blood type traits (8 items), 0.529 for the BFS (60-item Big Five personality test), and 0.501 for the TIPI-J (10-item Big Five personality test). In a comparison of the Big Five personality tests, the BFS, which has a larger number of items, was more accurate.

## 4. Discussions

### 4.1. Consistency with Previous Studies

As for differences in relation to blood type, none of the TIPI-J’s personality factors and only one BFS factor were statistically significant (Tables 2–3). Conversely, all 8 blood type questions were significant (Table 4). Furthermore, the majority of differences based on gender or age were more pronounced than those based on blood type (Figure 1). In the 8 blood trait items, the scores for each blood type were highest for the corresponding blood type, except for “self-centered”; type AB scored 3.967, slightly greater than type B’s 3.938. However, this difference was not statistically significant. Therefore, it can be stated that virtually all of the blood type traits in this study confirmed each score of the corresponding blood type was the highest. This evidently illustrates the correlation between blood type and personality. These results suggest that, as anticipated, question items with smaller differences were eliminated, and that tests with more items were more likely to detect differences in blood type.

**Figure 1.**
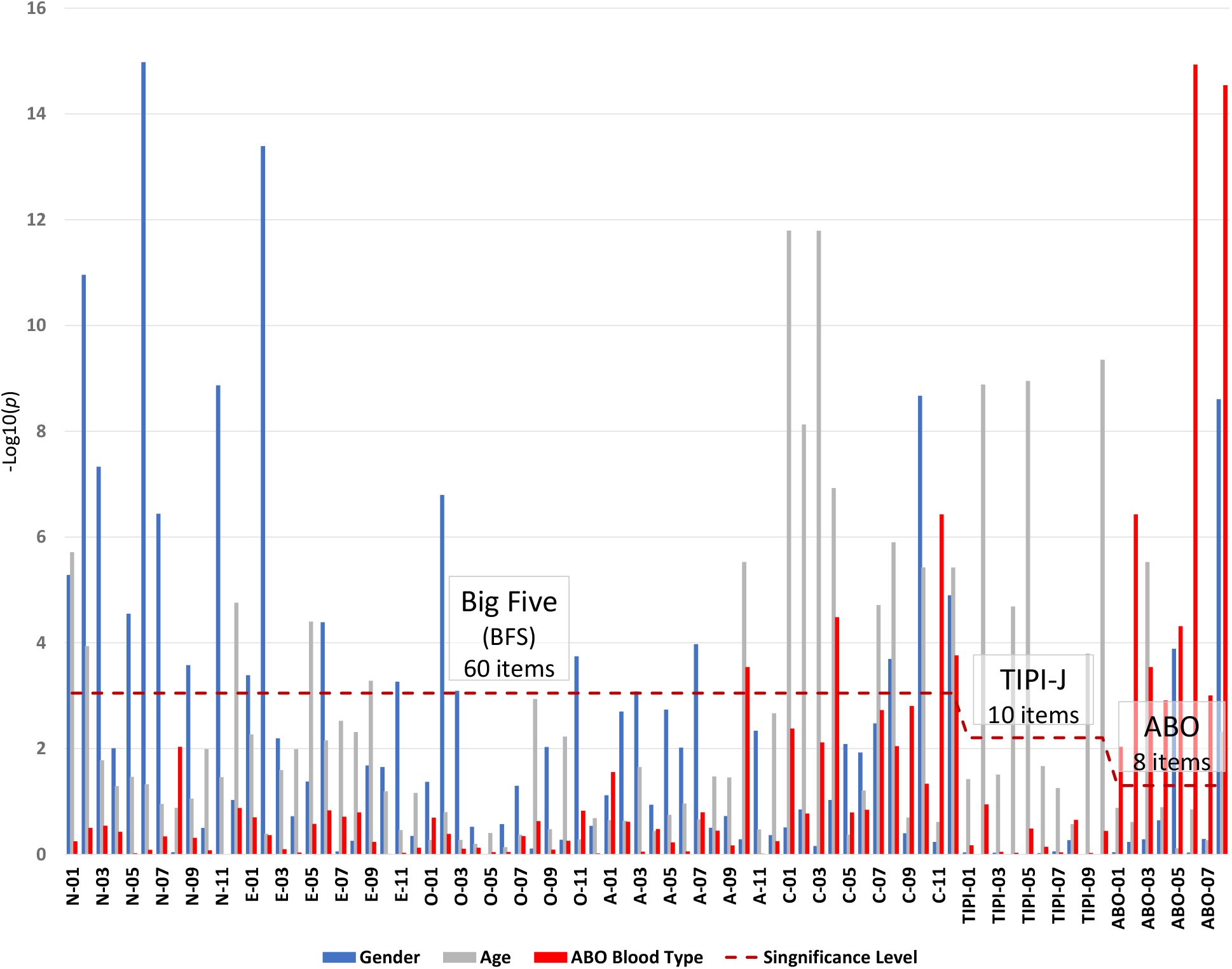
*p*-Values of Individual Items by Gender, Age and ABO Blood Type

It is generally acknowledged that the influence of blood type on personality is less pronounced than that of gender or age. Many prior studies have assumed that the effects of blood type are more significant than those of gender or age, leading to inadequate control of extraneous variables. It is likely that the effects of gender or age typically outweigh those of blood type, resulting in inconsistent results from the Big Five personality tests. In support of this, even though each of the 12 items comprising the BFS-N, BFS-A, and BFS-C contains 1 item for each blood type traits, only the BFS-C exhibited a statistically significant difference, while the BFS-N and BFS-C did not. In the future, more rigorous control of sample characteristics will be required. It seems that there exists a certain structure within studies utilizing the Big Five personality tests that makes it challenging to discern differences in blood type traits. As Lo et al. have demonstrated [5], it would be unsurprising for the coefficient of determination *R^2^* between a single gene and the Big Five personality test to be less than 0.4%, even analyzing over 260,000 individuals in total. These findings provide a potential explanation for the perplexing phenomena observed in previous studies, such as inconsistent differences and minuscule differences even with large sample sizes.

It is worth noting that some researchers contend that differences in personality by blood type are not genuine, but rather pseudo-correlations resulting from the “self-fulfillment phenomenon.” However, these researchers have conducted limited studies that directly examine data and demonstrate the self-fulfillment phenomenon. Conversely, studies that have directly investigated this have found differences as blood type traits, even in “groups with no knowledge of blood type traits” [32]. Therefore, the validity of the claim that all differences in personality stemming from blood type are caused by the “self-fulfilling phenomenon” is somewhat questionable.

### 4.2. Personality Sensitivity

The TIPI-J utilized in this investigation comprises a mere 10 question items for the 5 Big Five personality factors. To ensure the precision of the test with a relatively limited number of items, all of which are paired with opposite meaning items; it is assumed that the scores on each paired items for each factor are essentially negatively correlated. The results obtained from the sample of 2,887 utilized in this study were as follows, with all pairs displaying negative correlations (Table 6).

**Table 5.** Results of Blood Type Predictions Using AI

| Data | Weighted AUC | N | Algorithm |
| --- | --- | --- | --- |
| BFS (5 Factors with 60 Items) | 0.529 | 2887 | Voting Ensemble |
| TIPI-J (5 Factors with 10 Items) | 0.501 | 2887 | Voting Ensemble |
| Blood Type Traits (8 Items) | 0.593 | 2887 | Voting Ensemble |

**Table 6.**
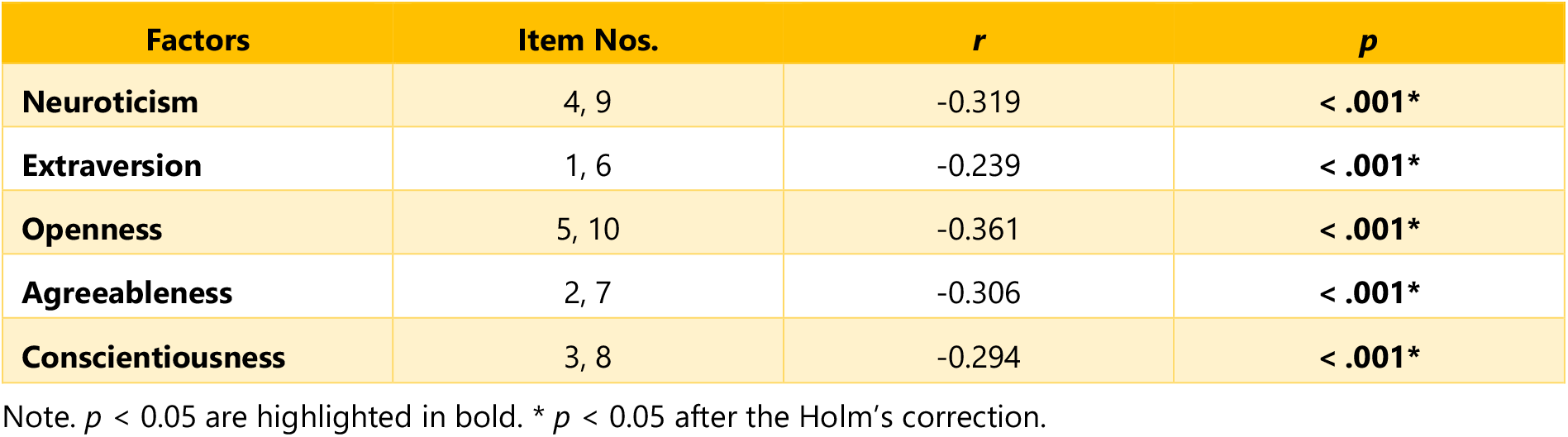
Correlation Coefficients between the TIPI-J’s Each Paired Items

However, certain prior studies [32] have revealed an anomalous phenomenon in which a correlation exists between greater levels of personality sensitivity and correspondingly greater scores. In other words, a “positive correlation” exists for certain question items that comprise these pairs. If this is not exceptional, it would significantly undermine the reliability of this particular personality test. To investigate this phenomenon, correlation coefficients were calculated between scores on the 10 TIPI-J question items and levels of personality sensitivity, utilizing a sample of 2,887 individuals. The results are illustrated in Table 7. Scores for personality sensitivity were determined by responses to the question “I am interested in my own and/or others personality” on a scale of 1 to 7 (with greater numbers indicating a greater interest).

**Table 7.**
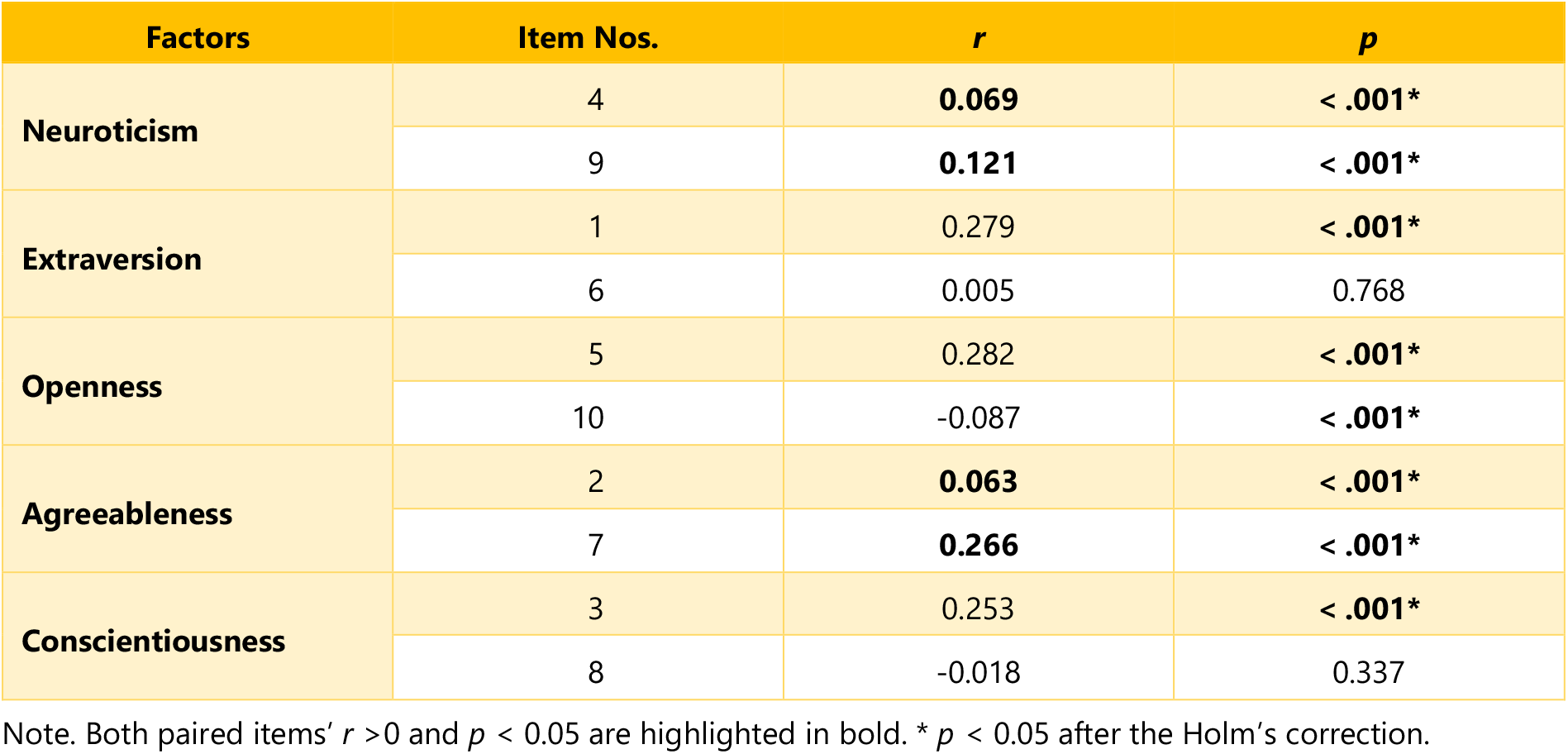
Correlation Coefficients between the TIPI-J Each Paired Items and Personality Sensitivity

Of the 5 pairs of question items, Neuroticism and Agreeableness were “positively correlated” with questions affirming personality sensitivity. Therefore, there were indeed pairs of question items that were essentially “opposite” in meaning, yet displayed much “the same” scores. The interpretation that appears most plausible is that, regardless of the specific questions asked, respondents with greater levels of personality sensitivity felt that they were more “matched” than others. This suggests that, even if scores on the Big Five personality test are the same, the personalities of the individuals in question may not necessarily be equivalent. This is close to the example of sensitivity testing mentioned above, in which there are individual differences in human perceptions and subjective and objective evaluations do not necessarily match.

By utilizing the same methodology as in Analysis 2, and utilizing data from only the 978 respondents who were deemed to possess high levels of personality sensitivity; those who affirmed both the “well knowledge” (respondents answered that they “know a lot” or “know some” with regards to blood type personality traits) and the “relationship between blood type and personality”, AUC increased from 0.593 to 0.628 (Table 8). This further supports the existence of an effect of personality sensitivity.

**Table 8.** Results of Blood Type Predictions Using AI

| Data | Weighted AUC | N | Algorithm |
| --- | --- | --- | --- |
| **BFS (5 Factors)** | 0.529 | 2887 | Voting Ensemble |
| **TIPI-J (5 Factors)** | 0.501 | 2887 | Voting Ensemble |
| **Blood Type Traits (8 Items)** | 0.593 | 2887 | Voting Ensemble |
| **Blood Type Traits (8 Items, Well-knowledge) \*** | 0.628 | 978 | Voting Ensemble |

\* Participants who answered both "they knew a lot (some) of blood type traits" and "blood type and personality were related a lot (somewhat)."

Additionally, the impact of personality sensitivity on scores for each item suggests that the influence of personality sensitivity was more pronounced than that of gender or age (Figure 2 and Table 9).

**Figure 2.**
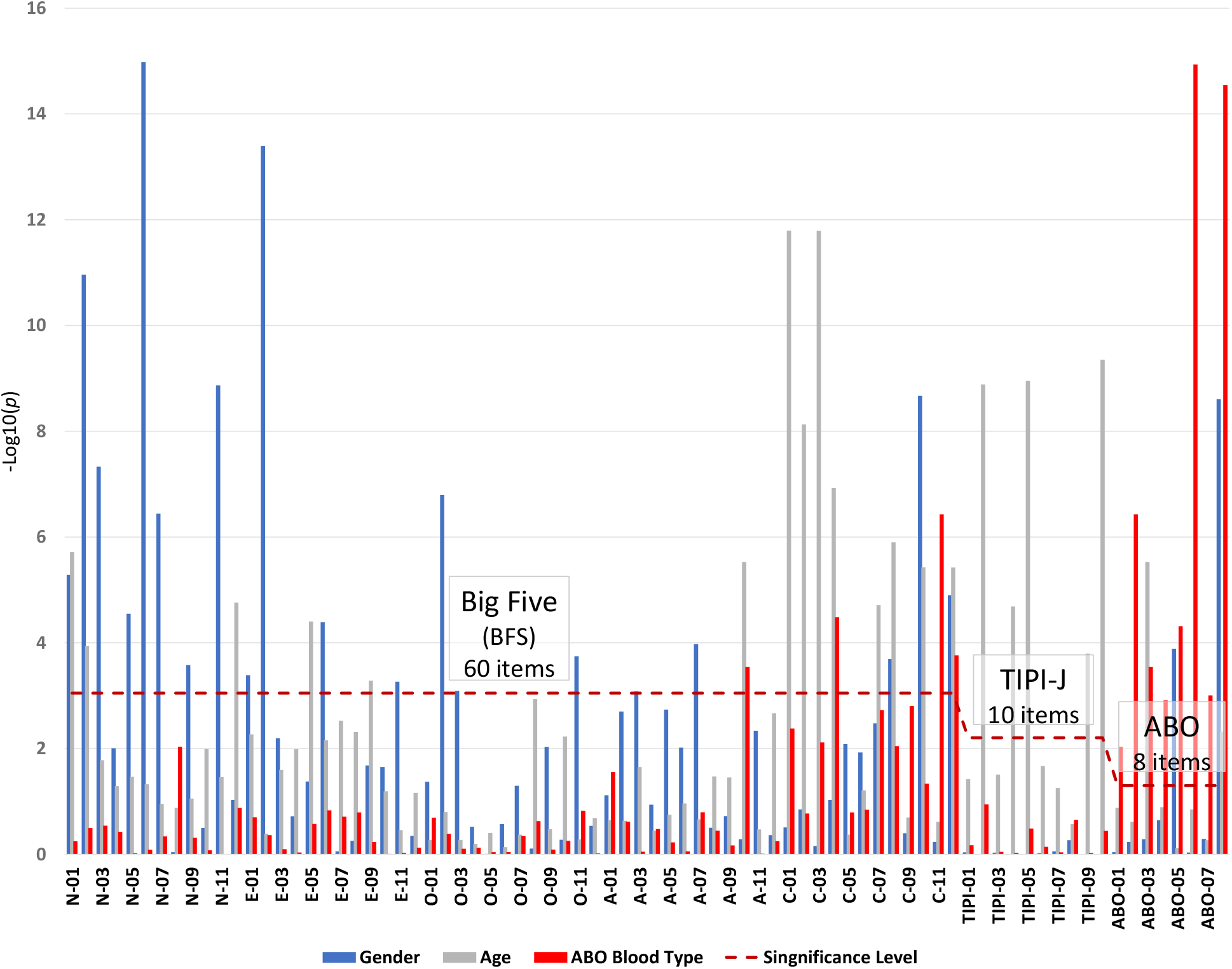
*p*-Values of Individual Items by Gender, Age and Personality Sensitivity

**Table 9.** Maximum *η^2^*-Values of ANOVA by Grouping Variables

| Data | Personality Sensitivity |  | Gender |  | Age |  |
| --- | --- | --- | --- | --- | --- | --- |
| | $\eta^2$ (Factor/Type) | $p$ | $\eta^2$ (Factor/Type) | $p$ | $\eta^2$ (Factor/Type) | $p$ |
| BFS | 0.126 (O) | 1.01e-80 | 0.008 (N) | 1.17e-06 | 0.047 (C) | 3.48e-13 |
| TIPI-J | 0.063 (O) | 6.37e-38 | 0.008 (N) | 7.56e-07 | 0.036 (C) | 4.57e-08 |
| Blood Type Traits | 0.042 (O) | 2.72e-24 | 0.019 (O) | 5.52e-14 | 0.022 (B) | 6.64e-03 |

Note. Each factor or type in the parenthesis was selected by those that showed the maximum value.

These may offer valuable insights regarding the interpretation of personality test scores, including the potential for Simpson’s paradox to be present. However, research on personality sensitivity remains limited [33–34]. Further studies in this area are eagerly anticipated.

### 4.3. Results of the Non-Big Five Personality Test

In addition to the Big Five Personality test, the relationship between blood type and personality has been investigated in various studies. One notable example is the study by Tsuchimine et al. [18], which had raw data that were publicly available; the data can be reanalyzed using AI and other methods. The Tridimensional Character Inventory (TCI) was employed in this study.

The TCI, a top-down personality model, is often used to examine genetic dispositions [35–36]. This personality test built a model for temperament with a physiological basis in the background. The test consists of 240 items using a yes-no scale rating.

Cloninger hypothesized that personality consists of traits that are hereditary and stable throughout life, and traits mature throughout life under the influence of socio-cultural environments. The TCI consists of 7 dimensions, including 4 temperament dimensions (Novelty Seeking, Harm Avoidance, Reward Dependence, and Persistence) and 3 character dimensions (Self-directedness, Cooperativeness and Self-transcendence). Three of the temperament dimensions have been hypothesized to be associated with monoamine neurotransmitters. Novelty seeking has been hypothesized to be associated with dopaminergic, harm avoidance with serotonergic, and reward dependence with noradrenergic.

In the study conducted by Tsuchimine et al., DNA analysis was utilized to ascertain the ABO blood types of the participants, comprising both genotype (AA, AO, BB, BO, OO, AB) and phenotype (A, B, O, AB). As phenotypes were employed in this study, the prediction target was set to be congruent with the phenotype. The publicly accessible data of 1,472 individuals did not comprise of gender or age, hence they were not included in the training data. The data incorporated genotypes (CC, CT, TT) of dopamine beta hydroxylase (DBH), a gene known to impact dopamine activity. Two groups of training data were formulated, one that included this gene and one that did not. To assess whether the disparities in blood types become progressively smaller as the number of personality factors decreases, supplementary data including DBH were also generated for the training data utilizing the 25 subscales of the TCI. Out of the total data, 1,427 individuals were employed, excluding the 45 individuals with missing values. The ABO phenotypes were predicted from the aforementioned 7 dimension and 25 subscales scores, by utilizing the same methodology as in Analysis 2. The results are depicted in Table 10. This appears to support our hypothesis that 1) multiple genes affect human personality, 2) a single gene affects a limited number of traits, and 3) in the personality factors, differences that appeared in single items become smaller or eliminated.

**Table 10.** Results of Blood Type Predictions Using AI including the TCI

| Data | Weighted AUC | Accuracy | N | Algorithm |
| --- | --- | --- | --- | --- |
| **BFS (5 Factors)** | 0.529 | 0.379 | 2887 | Voting Ensemble |
| **TIPI-J (5 Factors)** | 0.501 | 0.317 | 2887 | Voting Ensemble |
| **TCI (7 Dimensions) excluding DBH** | 0.536 | 0.367 | 1427 | Voting Ensemble |
| **TCI (7 Dimensions) including DBH** | 0.549 | 0.376 | 1427 | Voting Ensemble |
| **TCI (25 Subscales) including DBH** | 0.557 | 0.370 | 1427 | Voting Ensemble |
| **Blood Type Traits (8 Items)** | 0.593 | 0.390 | 2887 | Voting Ensemble |
| **Blood Type Traits (8 Items, Well-knowledge) \*** | 0.628 | 0.429 | 978 | Voting Ensemble |

\* Participants who answered both "they knew a lot (some) of blood type traits" and "blood type and personality were related a lot (somewhat)."

In this study, of the 3 groups examined with sample sizes of 2,887, 1,427, and 978 individuals (Table 10), the greatest number of individuals with type A were 1,091 (37.8%), 537 (37.6%), and 368 (37.6%), respectively. In other words, only the blood type traits clearly surpass the predictive accuracy of determining all as type A. This may imply certain limitations of contemporary personality tests.

In addition to personality tests, many academic researchers have analyzed the linkage between blood type and profession; notable differences were observed, particularly among politicians and athletes [8]. For instance, Ohmura, Ukitani, and Fujita, a team of Japanese psychologists, stated that “many Japanese prime ministers were type O (with a statistically significant level of *p* < 0.05)” [37]. A 2017 Italian study found that individuals with type O are predisposed to have superior athletic performance, particularly in long-distance running [38]. Furthermore, a 2009 Serbian study observed that the frequency of type O was higher among elite water polo players than other blood types [39]. These results also imply the potential relationship between blood type and personality or constitution.

### 4.4. Interpretation of Contradictory Phenomena

Assuming that the prior elucidations are adequate, it would be feasible to explicate the data on “blood type and personality,” which are commonly believed to be inconsistent, in a nearly unified manner.

For example:

1. In a study conducted in South Korea, there was no discernible variation in the Big Five personality test results of 200 participants. However, it was noteworthy that certain individual items on the Big Five personality test exhibited statistically significant differences, which mirrored certain traits associated with blood type. [12, 40]. Our study confirmed the reproducibility of this phenomenon.
2. In a study predicated on a comprehensive survey analyzing data from Japan and the United States on a population of over 10,000, virtually no differences by blood type were discovered. This is because the survey employed question items only pertaining to life and finances and did not comprise question items regarding personality [41].
3. Blood type traits have tended to remain constant over the several ten years, according to a periodic large-scale opinion survey (several thousand respondents annually, approximately 200,000) that comprises question items on personality traits [42–44].
4. The Big Five personality tests yielded equivocal outcomes or no differences due to the varying gender and age groups, which have a more pronounced impact on personality than blood type.

Some researchers have also cast doubt on whether the ABO blood type gene, a single gene, truly has a substantial impact on personality. However, recent studies have revealed that a single gene influences hundreds of other genes. For example, the gene BAZ1B, which is believed to be associated with human “self-domestication,” was found to affect the activity of 448 other genes [45]. This gene, BAZ1B, is crucial for neural crest and facial development and may be involved in self-domestication. Thus, there may be no particular quandary in assuming that ABO blood type genes influence personality.

## 5. Conclusion

The expression of traits by a singular gene is significantly limited. Nevertheless, a discernible correlation was observed between blood type and self-reported personality traits in various individual question items, which corresponded to traits that had previously been cited. Traditional personality tests, such as the Big Five test, usually employ a personality factor comprised of multiple items. Blood type does not exert a unique influence on an individual’s personality. On the contrary, various factors, such as gender and age, have a complex and notable impact. This precludes the attainment of consistent results without accounting for these nonlinear interactions. Traditional personality tests, such as the Big Five test, usually employ a personality factor comprised of multiple items. In this case, the effect of blood type is variable, which can lead to inconsistent results unless the gender, age or other conditions of the sample are properly controlled. The above explanation is closely aligned with the theory of personality psychology as well as other genetic effects phenomena. Our findings provide a novel, albeit hypothetical, framework for how genes impact human traits.

Meanwhile, the sample in this study was confined to Japanese populations only, the AI training data was of a small sample size, its use was experimental, and the interaction between gender and age is still unknown. Further research utilizing a larger, more global dataset is required to fully grasp the implications and to enhance methodologies.

## Acknowledgements

The author expresses appreciation to Chieko Ichikawa, Director of the Human Sciences ABO Center, for her assistance, as well as to Fred Wong, co-founder of AI Hong Kong Limited, for his guidance in utilizing AI. The author also extends gratitude to Professor Qinglai Meng of Oregon State University for his feedback and recommendations, and to Shoko Tsuchimine and her research team for providing the data in a format amenable to reanalysis.

## Supplemental File

S1 Dataset. Raw data. (XLSX)

## Appendix A

**Questionnaire in English (translated)**

Participants: 3,200 people

(Japanese males and females between the ages of 20 and 59, gender and age were equally allocated)

**Q1 – Q3.** BFS items (Appendix B)

**Q4.** ABO trait and personality sensitivity items

Please answer the following items on your personality.

Answer options

It does not fit me at all <- 4: Cannot say either way -> 7: It fits me well

Question items

01. Nervous (Type A, BFS N-08)
02. Meticulous (Type A, BFS C-11)
03. Self-centered (Type B, BFS A-10)
04. Hard to be understood (Type AB)
05. Self-paced (Type B)
06. Dual personality (Type AB)
07. Big-hearted (Type O)
08. Laid back (Type O)
09. Interested in your own or others personality (Personality sensitivity)

**Q5.** TIPI-J items

**Q6.** Do you think blood type and personality are related? (Relation)

1: Not related
2: Somewhat related
3: Strongly Related
4: I don’t know

**Q7.** Do you know the traits and the compatibilities of blood types? (Knowledge)

1: I don’t know at all
2: I know a little
3: I know some
4: I know a lot

**Q8.** Lastly, please tell us your blood type.

1: Type A
2: Type B
3: Type O
4: Type AB
5: I don’t know

## Appendix B

**BFS question items in English (translated)**

Personality trait terms based on the items of ACL (Adjective Check List)

**<BSF-E>**

01 Talkative
02 Silent R
03 Cheerful
04 Extravert
05 Introvert R
06 Unsociable R
07 Sociable
08 Misanthropic R
09 Active
10 Without expressing one’s intention R
11 Aggressive
12 Sober R

**<BFS-N>**

01 Overthinking
02 Anxiety-prone
03 Worried
04 Easily concerned
05 Faint-hearted
06 Easily hurt
07 Easily upset
08 Nervous (Blood type A trait)
09 Uncomfortable R
10 Pessimistic
11 Tense
12 Melancholic

**<BFS-O>**

01 Creative
02 Versatile
03 Progressive
04 Insightful
05 Imaginative
06 Aesthetically astute
07 Quick-witted
08 Flexible
09 Wide-ranging interests
10 Curious
11 Independent
12 Quick understanding

**<BFS-C>**

01 Irresponsible
02 Careless
03 Lazy
04 happy-go-lucky
05 Slothful
06 Designing R
07 Unconcerned
08 Thoughtless
09 Diligent R
10 Unprincipled
11 Meticulous R (Blood type A trait)
12 Easily bored

**<BFS-A>**

01 Mild-mannered
02 Short-tempered R
03 Angry R
04 Generous
05 Kind
06 Conscientious
07 Cooperative
08 Thorny R
09 Temperamental R
10 Self-centered R (Blood type B trait)
11 Honest
12 Rebellious R
(R: Reverse item)

**<Source>**

Sayuri Wada

Construction of the Big Five Scales of personality trait terms and concurrent validity with NPI

The Japanese Journal of Psychology 1996, Vol. 67, No. 1, 61-67.

https://doi.org/10.4992/jjpsy.67.61

